# Prioritizing Complex Disease Genes from Heterogeneous Public Databases

**DOI:** 10.1101/2023.02.09.527562

**Authors:** Eric Gong, Jake Y. Chen

## Abstract

**Background:** Complex human diseases are defined not only by sophisticated patterns of genetic variants/mutations upstream but also by many interplaying genes, RNAs, and proteins downstream. Analyzing multiple genomic and functional genomic data types to determine a short list of genes or molecules of interest is a common task called “gene prioritization” in biology. There are many statistical, biological, and bioinformatic methods developed to perform gene prioritization tasks. However, little research has been conducted to examine the relationships among the technique used, merged/separate use of each data modality, the gene list’s network/pathway context, and various gene ranking/expansions.

**Methods:** We introduce a new analytical framework called “Gene Ranking and Iterative Prioritization based on Pathways” (GRIPP) to prioritize genes derived from different modalities. Multiple data sources, such as CBioPortal, PAGER, and COSMIC were used to compile the initial gene list. We used the PAGER software to expand the gene list based on biological pathways and the BEERE software to construct protein-protein interaction networks that include the gene list to rank order genes. We produced a final gene list for each data modality iteratively from an initial draft gene list, using glioblastoma multiform (GBM) as a case study.

**Conclusion:** We demonstrated that GBM gene lists obtained from three modalities (differential gene expressions, gene mutations, and copy number alterations) and several data sources could be iteratively expanded and ranked using GRIPP. While integrating various modalities of data can be useful to generate an integrated ranked gene list related to any specific disease, the integration may also decrease the overall significance of ranked genes derived from specific data modalities. Therefore, we recommend carefully sorting and integrating gene lists according to each modality, such as gene mutations, epigenetic controls, or differential expressions, to procure modality-specific biological insights into the prioritized genes.

## Introduction

Gene prioritization has been an important research topic due to the rapid accumulation of experimental genomics data and the challenge of interpreting them in various biological contexts [1, 2]. For example, querying the gene signature database MSigDB with the term “breast cancer” can retrieve over 2000 statistically significant genes [3]. Creating a single therapeutic solution targeting more than a few target genes/proteins would still be technically challenging for drug development. Similarly, while pathway biomarkers have been proposed to monitor disease prognosis[4], it would be costly to test for all of them in clinical practice. Therefore, gene prioritization is critical for the feasibility of biomarker studies, drug development, or disease mechanism determination [5, 6]. Many approaches have been developed to prioritize genes specific to human diseases. One single method of prioritizing genes is based on measured gene expression changes, e.g., by “fold change”, which may be arbitrary to the choice of threshold values and the fact that some genes may change transiently or subtly to produce profound biological effects downstream [7]. Another method is to use p-value or other statistical significance measures to sort genes that changed under different conditions [8, 9]. However, p-value analysis can depend on the choice of samples; therefore, the results may not be transferable to new samples due to inherent sample biases [10]. Literature citation-based and text-mining approaches have also been used to curate significant genes related to a biological condition [1, 11, 12]. However, results may be biased toward literature studies that are “popular” and not necessarily directly indicative of biological importance and significance [1, 8]. Network biology-based approaches, e.g., prioritizing genes based on gene network centrality measures, may overcome many inherent biases of pure statistical or literature-based gene prioritization techniques [13-15].

Network-based gene prioritization tools today are developed with inherent assumptions and limitations, therefore having to be applied in practice with caution. Most prioritization tools, for example, may not provide necessary statistical details regarding the generated network. Instead, they may only provide a binary label [16]. Similarly, many network tools directly create the network without consideration for data quality. Thus, the responsibility of avoiding small sample coverage and filtering out noisy data from the pipeline falls to the users. For example, tools such as MGOGP [11] or KOBAS-i [12] are limited by their inability to expand gene lists. Thus, the output of these tools is restricted entirely by the quality of the input data; insufficient input data will lead to low quality or, even worse, biased results [17]. While the BEERE software tool [18] can expand gene lists, it fails to consider pathways and other biological contexts. Correctly placing genes into a biological context can aid the determination of significant genes not found within the initial query list [19]. The ability to parameterize search results—possible in tools such as BEERE and WINNER [8]—is also critical, allowing users to fine-tune results to their specific needs. Finally, almost no network tools implement a continuous, iterative refinement process. Iterative processes can help optimize the biological significance of results from network tools. The potential of many network-based prioritization tools may be hidden if multiple iterations are not conducted.

Here we introduce a new analytical framework called “Gene Ranking and Iterative Prioritization based on Pathways” (**GRIPP**) to address many of the limitations of current network prioritization methods. This framework addresses network biology challenges: providing quantitative statistical values to reflect biological significance, shifting the burden of data quality control away from users, and implementing an iterative process. In this study, GBM is used to illustrate the proposed workflow. However, the process can be easily generalized to any disease. We obtained the initial disease gene list (candidate genes) from several modalities, i.e., gene expressions, mutations, and copy number alterations, from several major biological databases. We then describe how to iterate through a network-based gene prioritization software tool (BEERE) and pathway enrichment/expansion software tool (PAGER) [20] to create a high-quality rank-ordered disease gene list that is biologically relevant.

## Methods

The process of procuring a high-quality disease gene list has four steps: compiling candidate gene subsets, gene prioritization based on pathway and network context, evaluation through literature co-citations, and evaluation through modality-specific gene ranks. The network-based gene prioritization and gene-set expansion tasks are iteratively repeated (as seen in Fig. 1 below) to refine the initial gene list into a highly query-specific and biologically significant gene list. To demonstrate this technique, we will build a case study using glioblastoma, one of the most aggressive and widespread brain tumors in adults [21]. However, the proposed methodology can easily be generalized to all diseases. To freely access the processed data, results, and a detailed step-by-step description of the technique in this work, one can visit https://github.com/aimed-lab/gene-prioritization

**Figure 1.**
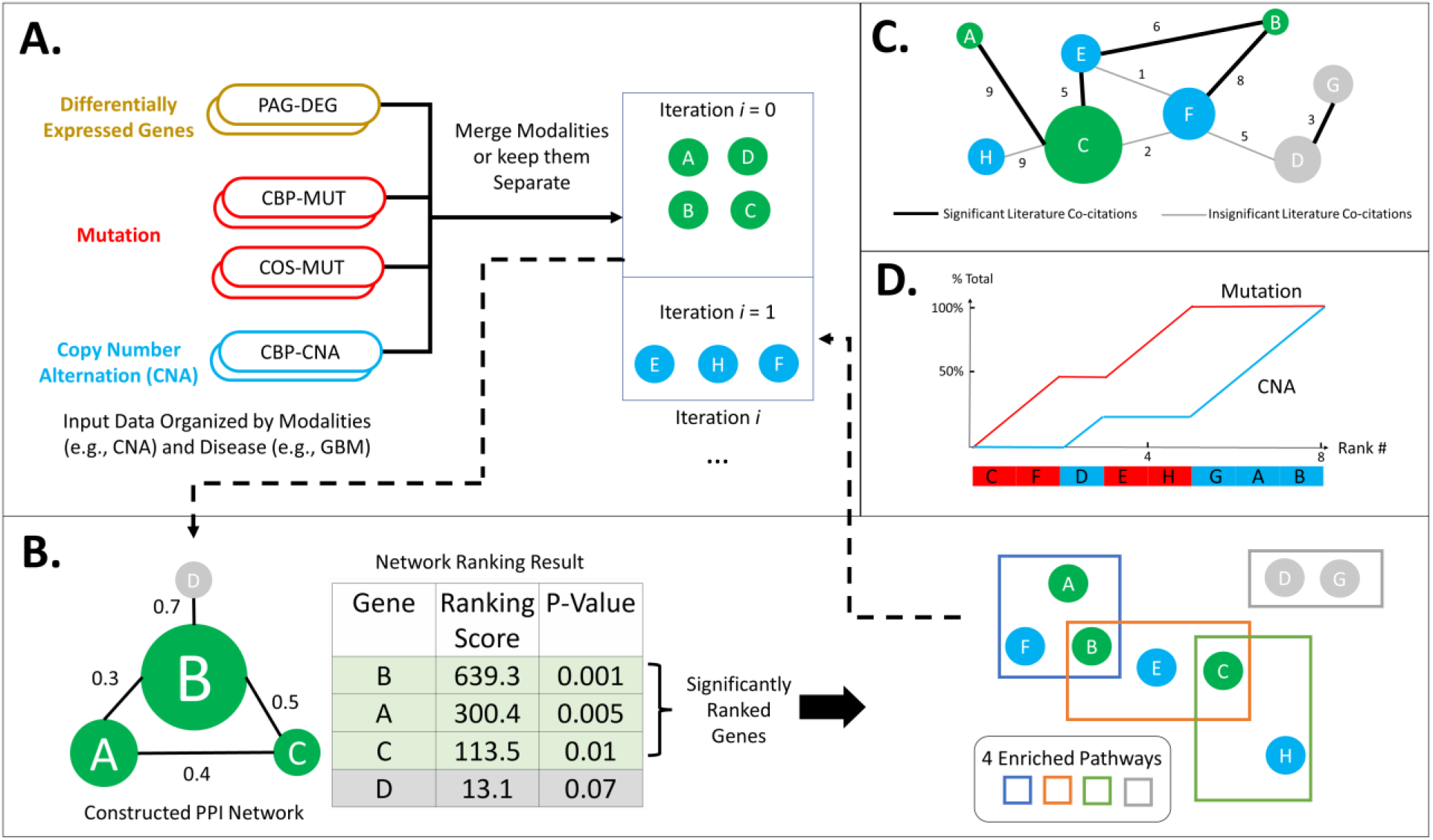
An overview of the Gene Ranking and Iterative Prioritization based on Pathways (GRIPP) framework. A) Compile seed gene lists (iteration *i* = 0), from different modalities such as Differentially Expressed Genes (DEG), Mutation (MUT), and Copy Number Alteration (CNA). They can be merged or kept separate as inputs to the gene prioritization process. B) First, construct a network, and pick any network gene prioritization software to generate significantly ranked genes; then, expand the resulting genes in their enriched pathway context to derive additional genes that will be added back to the next iteration (iteration *i =* 1, 2…) C) Overlay a literature co-citation network on top of the final PPI network involving all the resulting genes from all iterations. Node color indicates the batch of genes, whether significant (green or blue) or insignificant (grey), derived from the iterative process. D) A systematic evaluation of gene lists built from different modalities and their impact on gene rank orders.

### Compiling Candidate Gene Lists from Various Data Modalities

We created an initial disease gene list of five data subsets: PAG-DEG, CBP-CNA, CBP-MUT, COS-MUT, and COS-CTL. The COSMIC [22], CBioPortal [23], and PAGER [20] databases below are used to produce the initial disease gene list (candidate genes). Genes related to the query disease “Glioblastoma” were retrieved manually from databases, and a heuristic score was used to limit the number of genes retrieved for each data subset to less than 1200 for convenient downstream processing. Candidate genes from PAGER are differentially expressed genes and are represented by the symbol “**PAG-DEG**.” 663 PAG-DEG genes were heuristically selected with the PAGER nCoCo score, which signifies the biological relatedness of the gene sets contained within PAGER. Candidate genes from CBioPortal are copy number alteration genes and mutated genes and are represented separately as “**CBP-CNA**” and “**CBP-MUT**”, respectively. 653 CBP-MUT genes and 391 CBP-CNA genes were heuristically selected based on the frequency of copy number alteration and mutation occurrence, respectively. Candidate genes from COSMIC are mutated genes known to have a relation to the query disease and non-mutated control genes known to have no relation to the query disease and are represented separately as “**COS-MUT**” and “**COS-CTL**” respectively. 570 COS-MUT genes and 439 COS-CTL genes were heuristically selected based on the frequency of mutation, and lack of mutation frequency, respectively. To focus on only the highest-quality genes for downstream processing, BEERE was used to determine genes with significant p-value (p < 0.05). Here we differentiate between two combined gene lists. The first gene list is created from all five data subsets, is used to test the efficacy of BEERE gene prioritization, and is called ALL-CTL. The second gene list created from all data subsets except COS-CTL, will be used in the gene prioritization and refinement process and called ALL-EXP.

### Iterative Ranking and Expansion of Genes Based on their Network and Pathway Context

We refine the disease gene list using an iterative process involving a gene expansion and gene prioritization tool. Although currently existing tools are utilized, a unique pipeline is developed to coordinate these two tools to yield biologically significant results. We begin the iterative prioritization and expansion process by using BEERE to prioritize the disease gene list, determining genes with significant p-value (p < 0.05). BEERE is a user-friendly implementation of the WINNER software [8, 18]. Although BEERE only outputs one p-value per gene [18], it is sufficient for the prioritization methods employed within this methodology. We use Prioritization to select the top genes amongst the candidate and expand genes for further processing, yielding a more specific, biologically significant gene list. BEERE allows for the configuration of a direct p-value cut-off and direct access to a protein-protein interaction database with quality control measures. Thus, it is easy to parameterize the Prioritization process, sensitively controlling the size of the candidate or processed gene list, and as a result, the scope of the study.

Next, we query the prioritized gene list in PAGER to determine the related “pathway type” PAGs. The top upstream PAGs for each pathway PAG are then retrieved. The genes contained within each of the top upstream PAGs are then used to expand and enrich the gene list. The same heuristic score from the candidate gene compilation step – considering mutation frequency of the gene, frequency of overlap between different data sets, and the PAGER nCoCo score – is used to limit the number of expanded genes for convenient downstream processing before the expanded genes are combined with the original disease gene list. The gene enrichment step allows for significant genes not included in the initial candidate gene list to be added. This increases the overall quality of the gene list, and also aids in mitigating biases potentially present in the initial candidate gene list. The newly formed list is once again prioritized in BEERE, beginning another iteration of Prioritization, expansion, and refinement. The iteration is continued until less than 1% of the total expanded genes found through PAGER are not already present in the candidate gene list. In other words, the intersection of the candidate gene list and the expanded genes contains at least 99% of the expanded genes. The final high-quality gene list created from numerous iterations will be called ALL-FNL.

### Evaluation of Prioritized Genes in the Literature Co-citation Network

To assess the effectiveness of the methodology, we performed an evaluation of the ALL-FNL genes. One criterion used to evaluate the workflow is the biological significance of the ALL-FNL genes. To determine the biological significance of the ALL-FNL genes, we construct a network of statistically significant literature co-citations. This co-citation network is used to create a literature-based gene ranking, which is then compared against the ALL-FNL ranking generated through the iterative Prioritization process. To approximate the number of co-citations between glioblastoma and each retrieved gene, the two terms “Glioblastoma” and the gene name were queried in PubMed.

### Evaluation of Prioritized Genes through Modality-specific Gene Ranks

The second main criterion to judge the methodology is its ability to remove insignificant genes. The distribution and enrichment of the control genes within the gene list must be determined to evaluate the methodology’s ability to remove genes that do not have biological significance. Specifically, the Gene Set Enrichment Analysis (GSEA) software is utilized [24]. The ALL-CTL gene set –which contains the COS-CTL gene set known to have no significance to glioblastoma – is passed through one iteration of the methodology, with no cut-off parameter in BEERE, and then inputted into GSEA to determine where the COS-CTL genes would be placed in the gene ranking relative to the rank of the other modalities, demonstrating whether or not the methodology can indeed filter out non-significant genes.

## Results

In **Figures 2 and 3**, we can observe that the expanded genes deduced from the pathway and network-based biological context are highly ranked alongside and share many interactions with the well-established Glioblastoma genes. From **Figure 3**, we see that literature popularity is not directly correlated with gene ranking. Thus, looking at both literature and experimental data when deriving biological insight is necessary. For example, while there are 611 PubMed co-citations between SRC and RAC1, there are only 17 co-citations between SUMO1 and TP53. Nonetheless, both of these co-citations are considered significant. Thus, the number of co-citations and the prevalence of co-citations are not directly related to the degree to which the co-citation is biologically significant.

**Figure 2.**
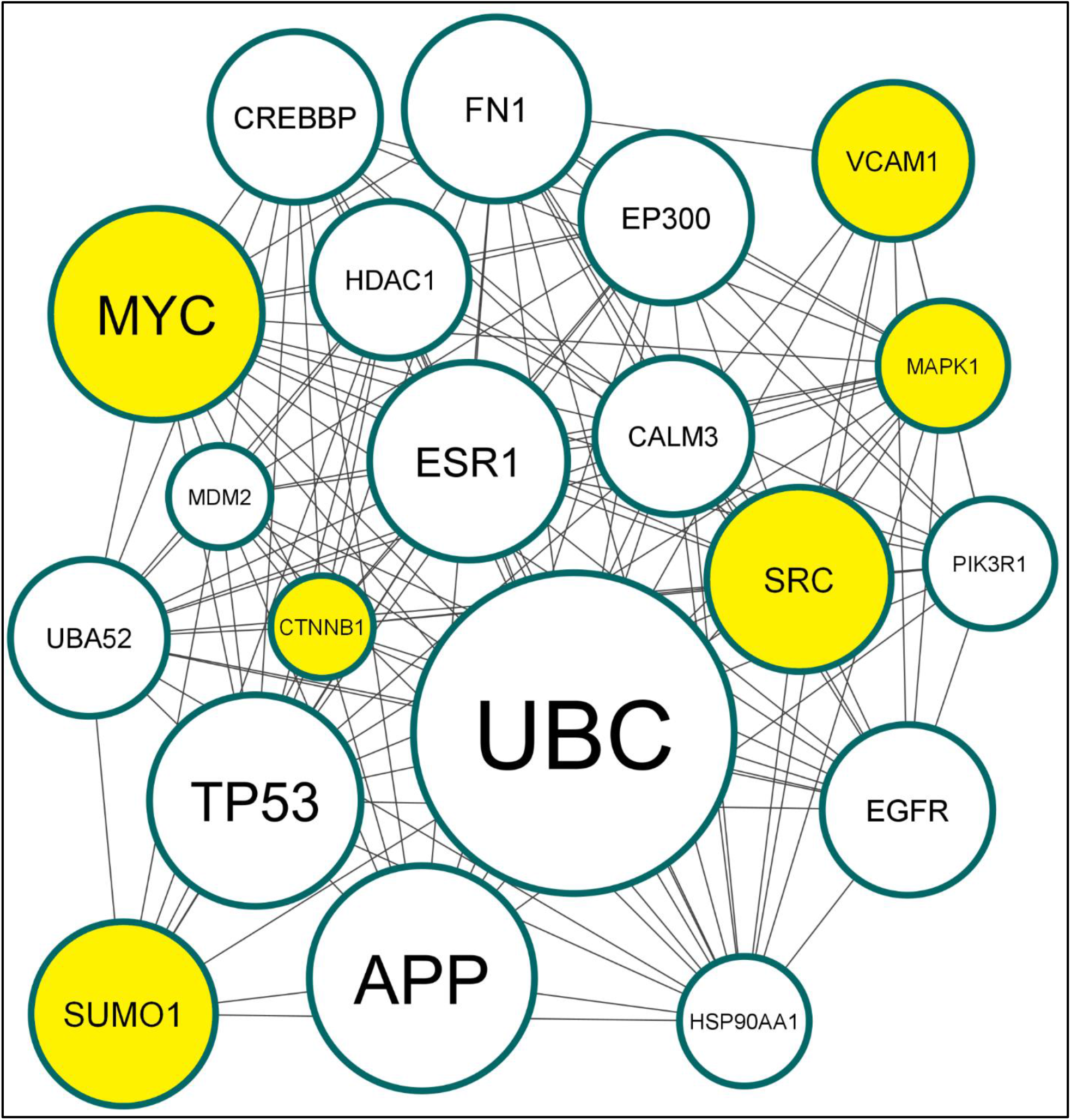
Gene refinement and prioritization using the query “Glioblastoma”. The gene network contains the top 20 ALL-FNL genes. Node size represents BEERE ranking score. Highlighted nodes are genes added through expansion based on biological pathway and network context.

**Figure 3.**
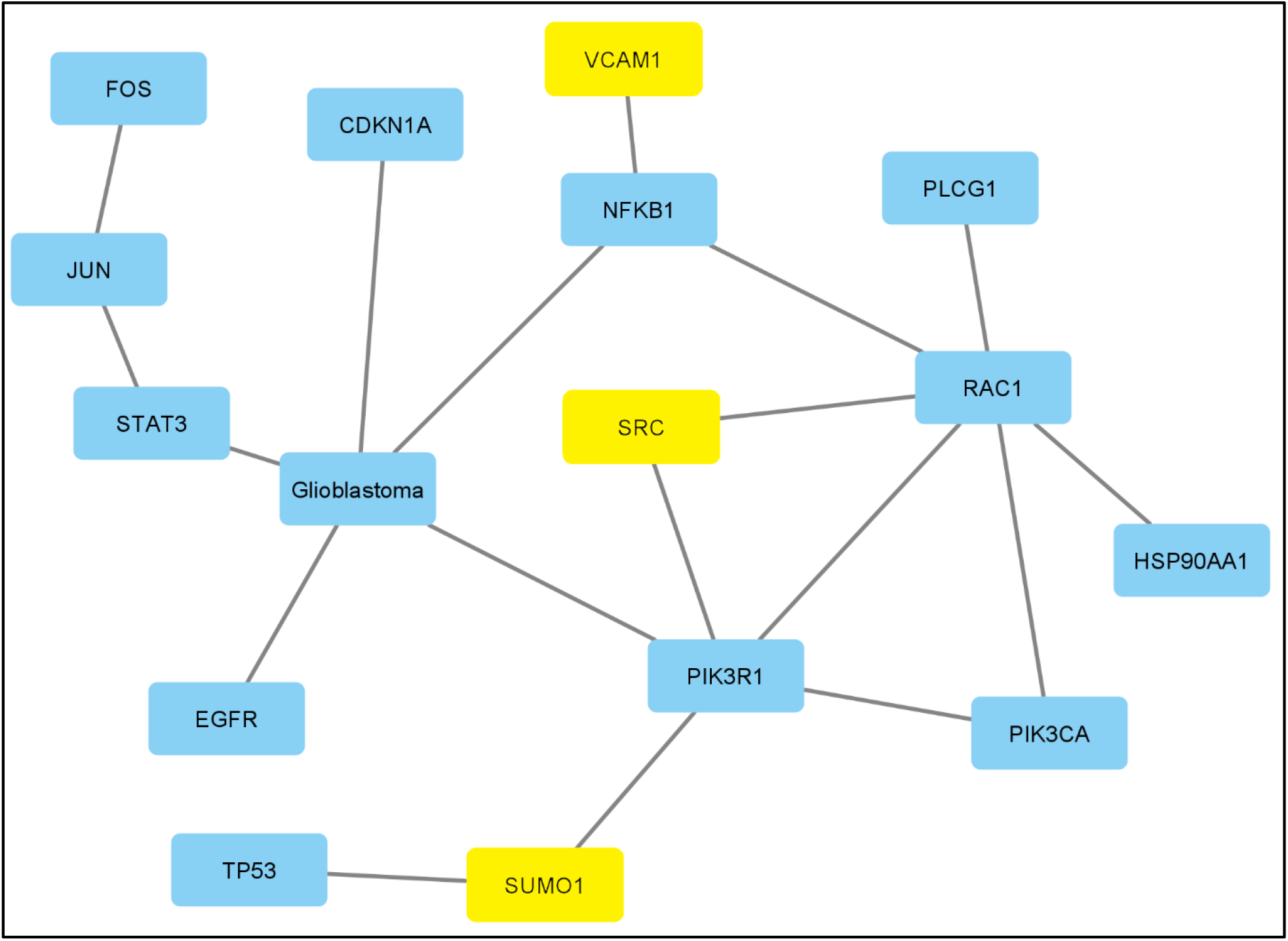
Statistically Significant Literature Co-Citation Network. The significant literature co-citation network is generated from the top 40 ALL-FNL genes. Highlighted nodes are genes added through the expansion based on a biological pathway and network context. Edges indicate statistically significant (p-value < 0.05) enriched co-citations in the literature of the two linked terms.

The COS-CTL genes are concentrated towards the lowest ranking, demonstrating the efficacy of BEERE in gene prioritization. However, we note that there is a preference for differentially expressed genes when compared to the mutated and CNA genes. Thus, there may be bias when integrating and combining multiple modalities.

## Conclusion

We determine three key conclusions. First, we demonstrate that an iterative network-based gene ranking on different -omic data modalities is biologically meaningful and that traditional network gene ranking can be refined through an iterative process. Our workflow case study using glioblastoma shows how users can combine or divide data according to modalities to generate combined or separate rankings. As seen in **Figure 2**, numerous genes discovered through the iterative Prioritization with pathway and network context are supported by statistically significant literature co-citations with glioblastoma. In addition, some genes added through the expansion process (not found in the initially compiled data subsets) are also significant in the literature, such as VCAM1, SRC, and SUMO1. As seen in **Suppl. Table 1**, the third-ranked gene, MYC, is an expanded gene not found in the initial data subsets, demonstrating the capacity of the iterative expansion process to determine significant genes not already present in the initial gene subsets. Second, gene rankings can reveal essential biology not reflected in the popularity of literature references. For example, there are only 22 co-citations between glioblastoma and the gene UBC (**Suppl. Table 1**).

Furthermore, UBC is not identified in BEERE’s statistically significant co-citation network, as seen in **Figure 2**. However, based on PAGER analysis, UBC is present in various biological pathways, such as the RAC1 pathway and ErbB signaling pathway, and is therefore closely linked to Glioblastoma genes established in the literature, such as MYC, TP53, EGFR, and MDM2. Third, we acknowledge that integrating datasets of the same modality from various different sources can yield biologically significant results. However, incorporating other modalities of data can cause ranking signals to be diluted. Therefore, to discover subtle signals, we recommend separating each modality, for example, by gene mutations, epigenetic controls, or differential expressions, to procure modality-specific biological insight. In **Suppl. Table 1**, both COS-MUT and CBP-MUT are data subsets containing mutated genes significant to glioblastoma. However, it is clear that each data subset contains different genes, and therefore both contribute towards the final gene list. In **Figure 4**, we show that although gene prioritization can concentrate the COS-CTL genes – known to have no relation to glioblastoma – near the lower end of the ranking, it is indeed the case that the differentially expressed genes PAG-DEG are diluting the signal of the other data subsets. This can also be seen in **Suppl. Table 1**, the number of PAG-DEG genes in the top 20 ALL-FNL exceeds the number of genes in the top 20 ALL-FNL ranking from the other data subsets.

**Figure 4.**
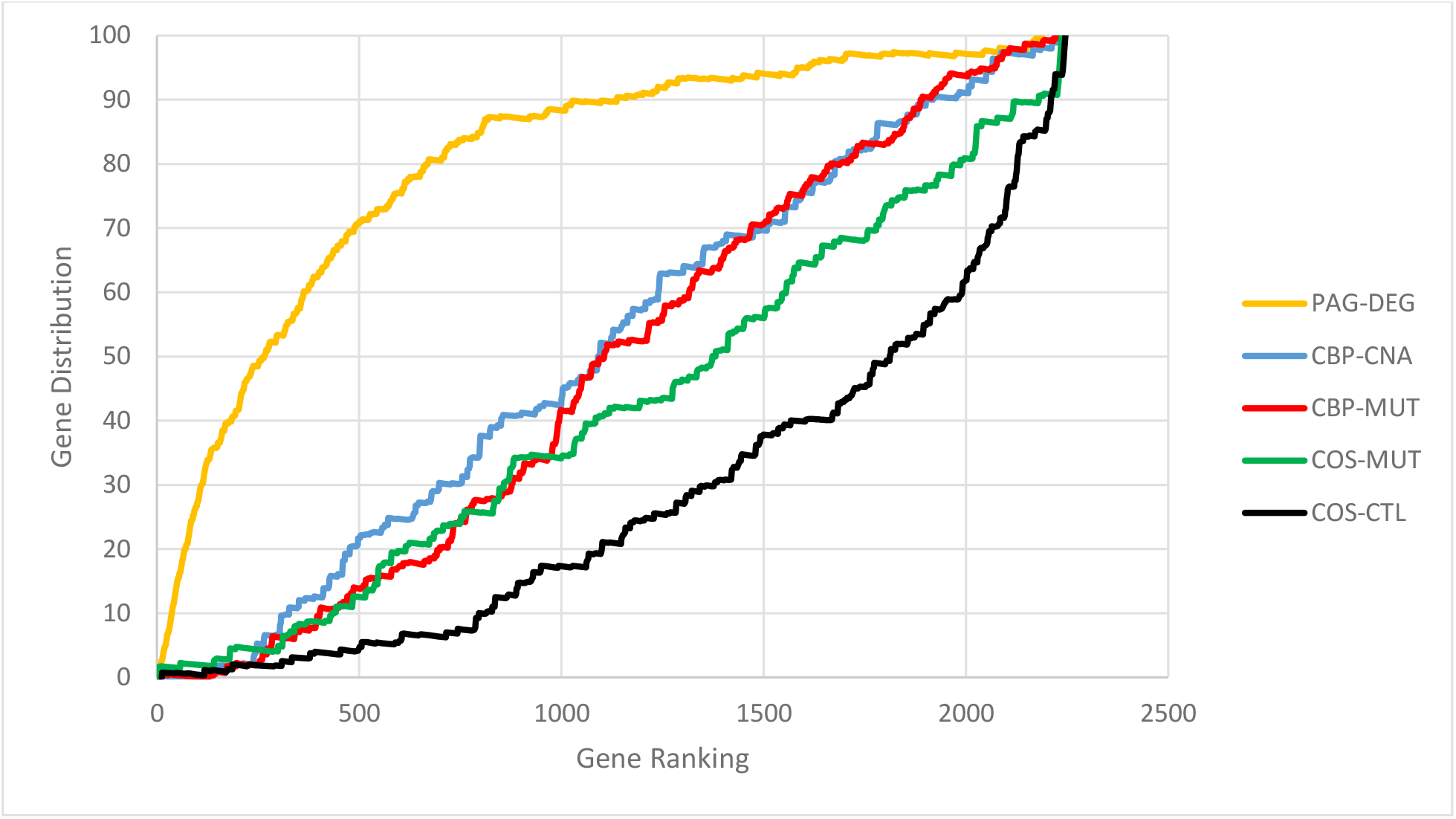
ALL-CTL gene distribution based on each underlying dataset analyzed by GSEA. The gene distribution is the percentage of genes found along the ranked list for each of the five data subsets, including the control data subset COS-CTL.

Using glioblastoma as a case study, we demonstrate how to extract biologically significant genes related to complex human diseases with GRIPP. To generalize our approach, one can compile a database containing refined gene lists for all human diseases, thus aiding researchers in future applications, such as prioritizing genes in drug discovery or choosing biomarker candidates. The generalization of our workflow could be accomplished by developing an API involving the PAGER Web APP in combination with WINNER – a streamlined implementation of BEERE – which will allow for iterative gene expansion and prioritization in a high-throughput fashion.

Depending on how a user wishes to use prioritized gene data, differing methods, and data modalities may be necessary. For example, if the Prioritization is used for sensitively discovering biomarkers, differentially expressed genes may be of more use. On the other hand, when discovering drug target candidates, modalities involving mutated genes may prove more useful.

## Acknowledgment

We thank the generous support of Dr. Jake Y. Chen, director of AI.MED lab at the University of Alabama at Birmingham (UAB), for mentoring this work in his spare time. JYC thanks the generous support of the UAB startup fund for this work. EG performed the data analysis using GSEA, drafted the manuscript, and conducted the data processing with PAGER and BEERE. JYC conceived the study, developed the data analysis plan, oversaw the implementation process by providing feedback throughout the research process, and revised the final manuscript before publication.

