## Supplemental Tables 1 and 2 for "Prioritizing Complex Disease Genes from Heterogeneous Public Databases"

### Supplemental Files

**Suppl. Table 1. Top 20 GBM network-ranked genes sorted by the overall ranking score and their ranks in each GBM data subsets.**

**Suppl. Table 2: A summary of the number of genes throughout each step of the GRIPP workflow based on network and pathway context.**

Note: Supplemental Excel Files containing the raw data from the data subsets, the iterative prioritization based on network and pathway context, and enrichment score results from GSEA can be found in our GitHub: https://github.com/aimed-lab/gene-prioritization

**Suppl. Table 1. Top 20 GBM network-ranked genes sorted by the overall ranking score and their ranks in each GBM data subsets.** Ranking scores were generated from the BEERE software (see methods) and the PubMed co-citation statistics was also listed.

| Gene Name | BEERE Ranking Score | Rank for each Gene Set | | | | | PubMed Co-citation with GBM |
|---|---|---|---|---|---|---|---|
| | | ALL-FNL | PAG-DEG | COS-MUT | CBP-MUT | CBP-CNA | |
| UBC | 626.9 | 1 | 1 | * | * | * | 22 |
| APP | 352.6 | 2 | 2 | * | * | * | 44 |
| MYC[+] | 204.9 | 3 | * | * | * | * | 496 |
| TP53 | 198.9 | 4 | * | 1 | * | 2 | 796 |
| ESR1 | 173.0 | 5 | * | 2 | * | * | 15 |
| FN1 | 155.3 | 6 | 3 | 3 | * | 1 | 29 |
| SRC[+] | 137.1 | 7 | * | * | * | * | 282 |
| SUMO1[+] | 133.9 | 8 | * | * | * | * | 7 |
| EP300 | 127.9 | 9 | * | * | * | 8 | 16 |
| EGFR | 127.4 | 10 | 5 | 4 | 1 | 3 | 2894 |
| CREBBP | 126.0 | 11 | * | 6 | * | 10 | 11 |
| VCAM1[+] | 124.8 | 12 | * | * | * | * | 32 |
| HDAC1 | 124.7 | 13 | * | * | * | 4 | 40 |
| CALM3 | 123.6 | 14 | 4 | * | * | * | 2 |
| UBA52 | 117.8 | 15 | 6 | * | * | * | 2 |
| HSP90AA1 | 110.8 | 16 | 8 | * | * | * | 5 |
| MAPK1[+] | 110.7 | 17 | * | * | * | * | 45 |
| PIK3R1 | 109.0 | 18 | * | 5 | 2 | 5 | 30 |
| CTNNB1[+] | 104.9 | 19 | * | * | * | * | 92 |
| MDM2 | 103.2 | 20 | 7 | * | 3 | * | 232 |

[+]Genes not present in the initial data subsets, and were added through the iterative prioritization based on pathway and network context.

**Suppl. Table 2: A summary of the number of genes throughout each step of the GRIPP workflow based on network and pathway context.**

| Description | Number of Genes |
|---|---|
| Number of initial candidate genes | 1857 |
| Significant genes ($p < 0.05$) after BEERE ranking | 242 |
| Genes discovered through expansion ($1^{st}$ iteration) | 591 |
| Significant expanded genes after BEERE prioritization ($1^{st}$ iteration) | 80 |
| Significant original genes after BEERE prioritization ($1^{st}$ iteration) | 214 |
| Total significant genes after $1^{st}$ iteration | 294 |
| Genes discovered through expansion ($2^{nd}$ iteration) | 591 |
| Significant expanded genes after BEERE prioritization ($2^{nd}$ iteration) | 290 |
| Significant original genes after BEERE prioritization ($2^{nd}$ iteration) | 13 |
| Total significant genes after $2^{nd}$ iteration | 303 |
